# Sex matters: the MouseX DW-ALLEN Atlas for mice diffusion-weighted MR imaging

**DOI:** 10.1101/2023.11.27.568800

**Authors:** Patricia Martínez-Tazo, Alexandra Santos, Mohamed Kotb Selim, Elena Espinós-Soler, Silvia De Santis

## Abstract

Overcoming sex bias in preclinical research requires not only including animals of both sexes in the experiments, but also developing proper tools to handle such data. Recent work revealed sensitivity of diffusion-weighted MRI to glia morphological changes in response to inflammatory stimuli, opening up exciting possibilities to characterize inflammation in a variety of preclinical models of pathologies, the great majority of them available in mice. However, there are limited resources dedicated to mouse imaging, like those required for the data processing and analysis. To fill this gap, we build a mouse MRI template of both structural and diffusion contrasts, with anatomical annotation according to the Allen Mouse Brain Atlas, the most detailed public resource for mouse brain investigation. To achieve a standardized resource, we use a large cohort of animals *in vivo*, and include animals of both sexes. To prove the utility of this resource to integrate imaging and molecular data, we demonstrate significant association between the mean diffusivity from MRI and gene expression-based glial counting. To demonstrate the need of equitable sex representation, we compared across sexes the warp fields needed to match a male-based template, and our template built with both sexes. Then, we use both templates for analysing mice imaging data obtained in animals of different ages, demonstrating that using a male-based template creates spurious significant sex effects, not present otherwise. All in all, our MouseX DW-ALLEN Atlas will be a widely useful resource getting us one step closer to equitable healthcare.

**Graphical abstract:** 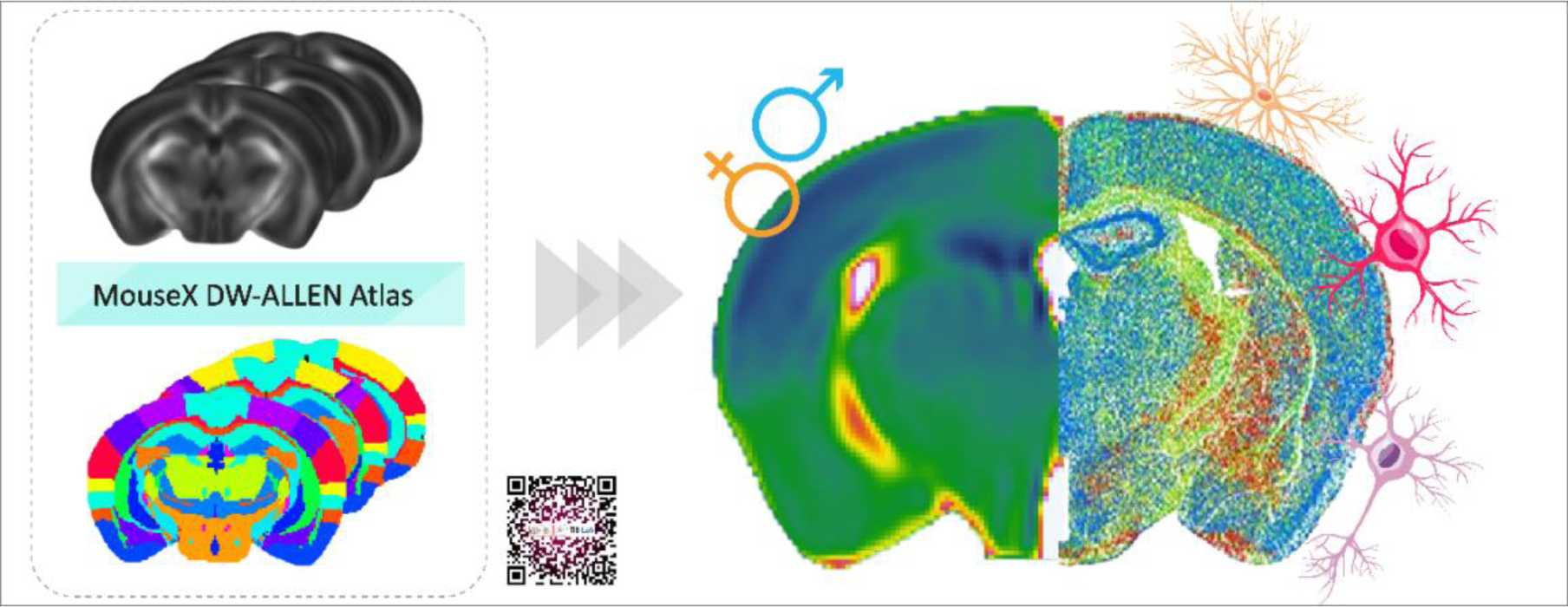

## Introduction

Sex bias in research leads to skewed findings and misconceptions about the true effects of treatments, drugs, or interventions [1]. This bias not only undermines the accuracy and reproducibility of research outcomes, but also hampers progress toward personalized and equitable healthcare. While efforts have been made to involve females in clinical studies, progress in this regard has been limited in the preclinical realm [2], [3], even though it lays the foundation for subsequent clinical trials.

Magnetic Resonance Imaging (MRI) holds a huge potential in preclinical research, by opening a window *in vivo* to brain microstructure which, complemented with a wide range of molecular techniques, can offer a unique translational characterization in a variety of animal models of pathologies. For example, our laboratory has recently proposed a strategy to reveal glia reactivity *in vivo* using diffusion-weighted MRI (DW-MRI), opening an unprecedented scenario to characterize microglia morphology in health and disease [4]. Indeed, the nervous and immune systems are in constant bidirectional interaction via chemical and cell-to-cell interactions [5], [6], with immune system playing both protective and detrimental role [7]. Therefore, characterization of brain parenchyma inflammation can serve as a crucial early biomarker in neurodegenerative diseases.

Importantly, sexual dimorphism has a prominent role in neuroimmunity [8]. Driven by sex steroids, gene expression encompassing immune cell maturation differs in male and female microglia [9], resulting in phenotypic and functional dimorphism, that is maintained in adulthood leading to different region-wise microglia density and proteomic profile [10]. In addition, glial cells from male and females have intrinsic sex differences, as evident in tissue cultures [11]. Interestingly, DW-MRI was used to highlight significant interactions between age and gender in humans, showing that microstructural senescence starts earlier in males [12], but the neurobiological substrate of these differences remains untapped. Characterizing sexual dimorphism in animal models could help elucidate this important issue.

While the mouse is the animal model of choice for neurodegenerative diseases thanks to its suitability for genome modifications, traditionally the rat has been preferred for MR imaging due to larger head size. As a result, currently available analysis resources are mostly suitable for rats. There is a pressing need to develop tools specific for mice, like a brain standard atlas for registration and region annotation. In this context, the Allen Mouse Brain Atlas (AMBA) is an open-source large-scale database collecting info from several fields within a common 3D coordinate framework (http://www.brain-map.org). A large *in situ* hybridization study has allowed the creation of a highly standardized genome-wide atlas of gene expression aligned with the AMBA annotation [13]. However, while the AMBA coordinate framework has been adapted for many computational tools and imaging techniques, such as automatic histological slice compartmentalization [14], spatial transcriptomics [15], [16], and two-photon serial tomography [17], there is no AMBA adapted for *in vivo* diffusion-MRI analysis yet, and built with animals of both sexes.

Here, our aim is to provide a DW-MRI mouse reference atlas under the consensus AMBA annotation, and including animals of both sexes. With this idea in mind, we scanned a large cohort of animals *in vivo*. We generated a template with three different contrasts, one relaxometry based (T2-weighted), and two diffusion-based contrasts (Fractional anisotropy, FA; and Mean diffusivity, MD) with high resolution. As sexual dimorphism in C57BL/6J mice has previously been reported [18], both female and male mice were included in the template generation. Then, we adjusted the AMBA annotation to the MRI template, generating the MouseX DW-ALLEN Atlas, which delineates a totality of 63 regions: 45 grey matter cortices and subcortical nuclei, 17 white matter tracts, and the ventricular system.

With this resource, we demonstrated the potential that the MouseX DW-ALLEN Atlas holds to merge imaging and molecular information by testing the correlation between MD, an imaging marker sensitive – albeit nonspecific - to microglia density [19], and gene expression-based glial counting from the AMBA delineated by Erö et al [20]. In addition, we demonstrated the need to use a brain template built with animals of both sexes by comparing the deformation fields that female brains undergo to fit a male-based template, as compared to the deformation to a template created with both sexes. Finally, we acquired a new mice dataset and investigated the impact of using a male-only template, demonstrating that using a male-based template creates spurious significant sex effects, not present otherwise.

All in all, the MouseX DW-ALLEN Atlas is expected to have a large impact to facilitate *in-vivo* mouse brain DW-MRI and foster the inclusion of female animals in preclinical research, a much-needed step to overcome gender bias. All data, generated templates, and the MouseX DW-ALLEN Atlas are publicly available at (https://github.com/TIB-Lab/MouseX-Allen-Atlas).

## Methods

### Animal preparation

All animal experiments were approved by the Institutional Animal Care and Use Committee of the Instituto de Neurociencias de Alicante, Alicante, Spain, comply with the Spanish (law 32/2007) and European regulations (EU directive 86/609, EU decree 2001-486, and EU recommendation 2007/526/EC) and the ARRIVE recommendations. 60 C57BL/6J mice (30 females, 30 males) were housed in groups of 5 (females separated from males), with 12-hour/12-hour light/dark cycle, lights on at 8:00, at room temperature (23 ± 2°C) and with free access to food and water. Animals were scanned at 56-60 postnatal day. A second cohort of C57BL/6J mice (n=9, 4 females), control littermate of APP/PS1 mice used in another experiment, was included for a longitudinal experiment, and scanned at 90, 141 and 190 days post-natal.

### MRI experiments

Relaxometry (T2-weighted) and DW-MRI experiments were conducted on the first cohort of mice (n=60) on a 7-T scanner (Bruker, BioSpect 70/30, Ettlingen, Germany) using a receive-only phase array coil with integrated combiner and preamplifier in combination with an actively detuned transmit-only resonator. For high resolution T2-weighted (T2W) data, a RARE (Rapid Imaging with Refocused Echoes) sequence with a matrix size = 258 x 214, in-plane voxel size = 70 µm^2^, forty slices of 400 µm thickness, repetition time (TR) = 3394 ms, and echo time (TE) = 33.5 ms were used. Both sequences were acquired with 4 averages and 1 repetition. In a subset of these same animals (20 males and 20 females), DW-MRI data were acquired using a spin-echo EPI (Echo Planar Imaging diffusion) sequence, with 20 uniform distributed gradient directions, b = 1000 s/mm^2^, diffusion time 15 ms with four images without diffusion weight (b = 0, called b0), TR = 8000 ms, TE = 26 ms. Forty axial slices of 400 µm thickness were set up with field of view (FOV) = 18 x 15 mm, matrix size = 120 x 100 and in-plane voxel size = 150 µm^2^. Total scan time including animal positioning was around 1 hour. The second cohort of animals (n= 9, 4 females) was scanned with the same scanner and coil. For T2W data, a RARE sequence with a matrix size = 108 x 90, in-plane voxel size = 160 µ m^2^, sixteen slices of 800 µm thickness, TR = 3000 ms, TE = 7.7 ms were used, acquired with 4 averages and 1 repetition. DW-MRI data were acquired for the same cohort using a stimulated echo EPI sequence, with 20 uniform distributed gradient directions, diffusion time 15 ms, b = 1000 s/mm^2^, four b0s, TR = 8000 ms, and TE = 25 ms, acquired with 1 average and 1 repetition. The geometry was the same of the T2W acquisition.

### MRI analysis: Template generation

MRI raw data was processed using in-house Matlab (MATLAB R2021a) scripts as follows. The ANTs (Advanced Normalization Tools) [21] registration framework was used for all the image registrations. FSL BET (Brain Extraction Tool) [22] was used to remove the skull from the T2W and DW-MRI data, followed by manual correction. Using the first mice cohort, a T2W template was created by recursively co-registering, bias-correcting, and averaging single subject skull-stripped T2W images using the ANTs tool *buildtemplateparallel.sh* with seven iterations. The obtained warp transform files were used to compute the single-subject Jacobian determinant (using the ANTs tool *CreateJacobianDeterminantImage*). Skull-stripped DW-MRI data was nonlinearly registered to the skull-stripped T2W data to correct for EPI distortions, followed by correction of motion distortions by affine registration to the b0, free water elimination [23], and lastly denoised with patch2self, a self-supervised learning method [24]. Subsequently, the DW-MRI images were processed for Diffusion Tensor analysis employing a robust model fitting approach [25] using the toolbox ExploreDTI [26]. Mean diffusivity (MD) and fractional anisotropy (FA) maps were extracted for each subject and normalised to the T2W template by applying the transformation calculated using the T2W images. Finally, the normalised microstructural maps of the individual subjects were merged and averaged to obtain MD and FA contrasts. For white-matter voxel-wise analysis, the mean FA skeleton across all subjects was created from the Tract-Based Spatial Statistics (TBSS) function *tbss_3_postreg*, by using the generated FA template as input. An FA threshold of 0.2 was applied. The workflow is illustrated in Figure 1. To compare deformations to a both sexes template with deformations to a male-only template, the same procedure to generate the template was repeated including only male mice. For completeness, a female-only template was also generated.

**Figure 1.**
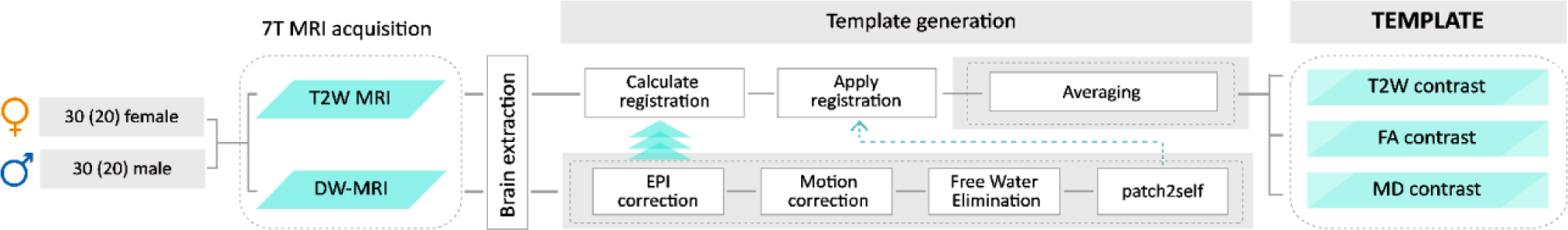
Workflow. Experimental workflow and processing steps employed for the template generation. Image acquisition of high-resolution T2W and DW-MRI sequences was performed on a total 60 mice (30 female) for T2W contrast, and 40 (20 female) for the diffusion-weighted contrasts. Image processing begins with skull removal. T2W images are used for template creation with the function *buildtemplateparallell*, which recursively calculates and then applies the registration to the common space. DW-MRI images undergo EPI distortion correction, followed by correction of motion distortions, free water elimination, and denoising with patch2self; finally DTI maps are extracted for FA (Fractional Anisotropy) and MD (Mean Diffusivity). The DTI maps are brought in the T2W template space and averaged to generate the FA and MD template contrasts.

### MRI Parcellation: AMBA CCFv3 reference atlas

The two-photon tomography-based annotation, the most advanced 3D mouse brain atlas with a detailed histology-based segmentation and delineation of 43 isocortical areas and their layers, 329 subcortical GM structures, 81 fiber tracts, and 8 ventricular structures per hemisphere [17], served as starting point to parcellate our MRI template. The AMBA CCFv3 annotation and template were downloaded from https://scalablebrainatlas.incf.org/mouse/ABA_v3, namely the “P56_Annotation_downsample2.nii.gz” and “P56_Atlas_downsample2.nii.gz” files. CCFv3 template was then non-linearly registered to the T2W template. An affine transformation with a neighbour cross-correlation metric was used, followed by a SyN transformation with Mattes metric. Subsequently, the affine and inverse warp transforms were applied to the AMBA CCFv3 annotation to bring it in the MRI space, using the nearest neighbour interpolation method. Then, all 40 serial slices were individually revised and corrected voxel-wise, following Allen Mouse Brain Coronal Atlas annotation (https://mouse.brain-map.org/static/atlas), to maximise anatomical correspondence, and, importantly, reduce the number of parcellated regions. In fact, due to the lower MRI template resolution as compared to the CCFv3 Template (40 μm^2^ compared to 10 μm^2^), the AMBA annotation was reduced to 63 ROIs, following anatomical hierarchy-guided region merging.

The first round of manual corrections, mostly focusing on grey matter annotation, was done using the T2W contrast as a reference; a second refinement using diffusion contrast (FA) as a reference focused instead on optimal white matter segmentation. The experimental workflow is illustrated in Figure 4A.

### Leveraging imaging and molecular data: correlation to Allen ISH data

To demonstrate the potential of sharing a common annotation between different neuroimaging techniques, we tested for significant associations between MRI data and molecular data from the Allen repository. The 3D Cell Atlas for the Mouse Brain delineated by Erö et al [20] is the first 3D atlas showing cell positions constructed algorithmically from whole brain cell-specific gene expression stains. It provides cell position, density and number of neurons, astrocytes, oligodendrocytes and microglia, in 737 grey matter brain regions defined as in the AMBA. The data from the 737 regions was merged to match the 45 grey matter regional delineation from our MouseX DW-ALLEN annotation; when our annotation comprised several regions of the AMBA, cell counts have been summed. As MD is an imaging marker sensitive to microglial density and reactivity profile [19], signal intensity measure from the MD contrast of the template was extracted region-wise, and Spearman correlation coefficient was calculated between MD measure and each individual cell type count. P-values were corrected for multiple comparisons across tested conditions according to false discovery rate approach [27].

### MRI statistics: sexual dimorphism

The voxel-wise comparison between the deformation fields in males and females to match the MouseX DW-ALLEN template, and its male-only version, was evaluated using the randomise threshold-free cluster enhancement function (from the FSL library [22]) on the Jacobian determinant [28]. A general linear model was used to test for sex effect, while correcting for multiple comparisons across voxels. Then, in the second cohort of animals scanned at three time points, the two templates (both sexes and male-only) were used to extract, for each animal and time, the average MD values in each atlas region. A general linear model with age and sex as factors was tested in each region of interest, and the p-value was adjusted for multiple correction using false discovery rate [27].

## Results

### The MouseX DW-ALLEN Atlas

We have generated a mouse MRI 3D brain template, with three contrasts, as detailed in Materials and Methods and following the workflow detailed in Figure 1: T2W anatomical contrast, and DW-based FA and MD. Representative sagittal, coronal and axial planes are reported in Figure 2. In order to facilitate execution of the Tract-Based Spatial Statistic, the standard statistical approach to test general linear models in white matter [29], we have also generated a white matter skeleton, as shown in Figure 3.

**Figure 2:**
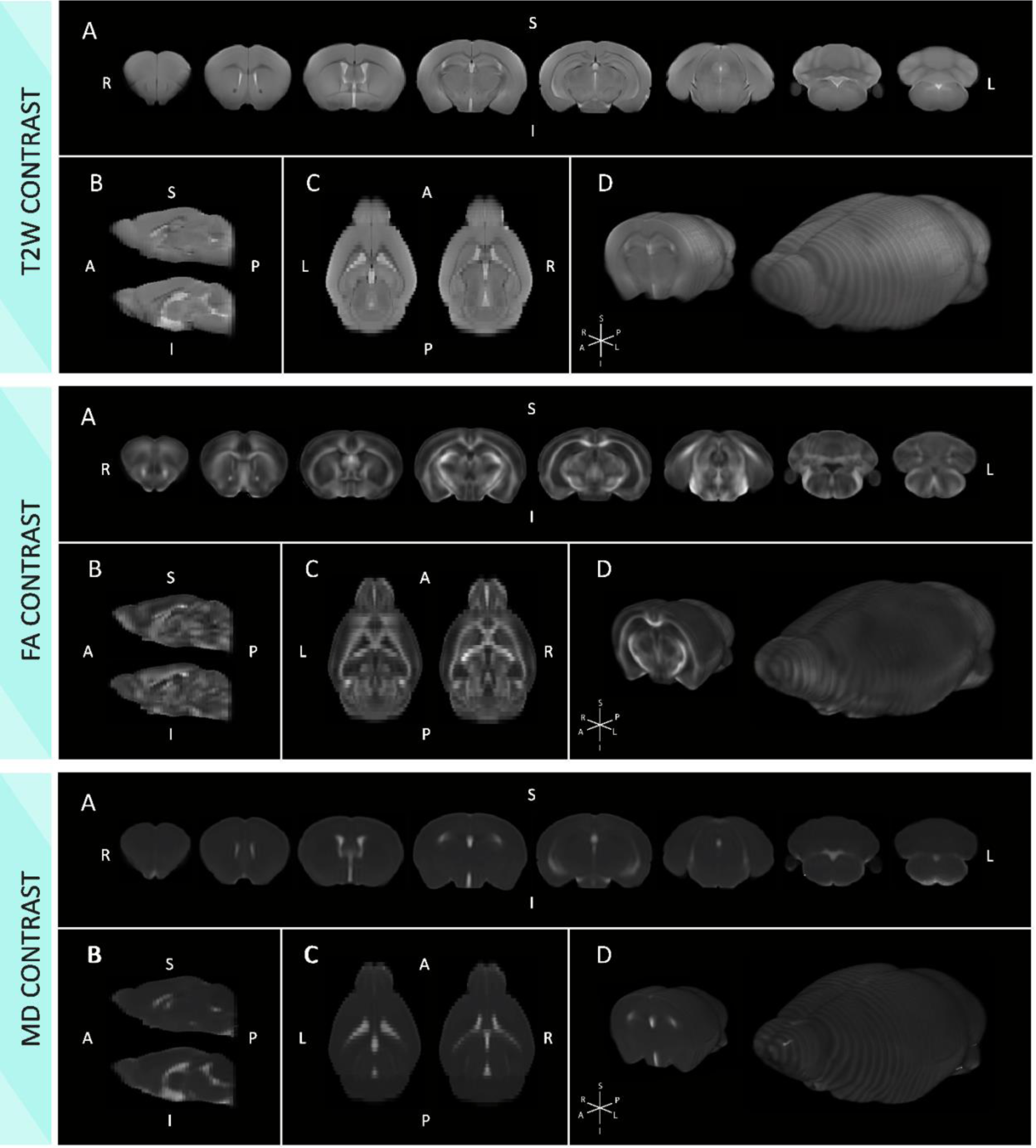
The MouseX DW-ALLEN multi-contrasts template. High resolution mouse anatomical templates for T2W contrast (top), FA contrast (middle) and MD contrast (bottom). A) coronal, B) sagittal and C) axial slices of the average template generated from 30 males and 30 females (20 each for FA and MD). D) three-dimensional rendering.

**Figure 3:**
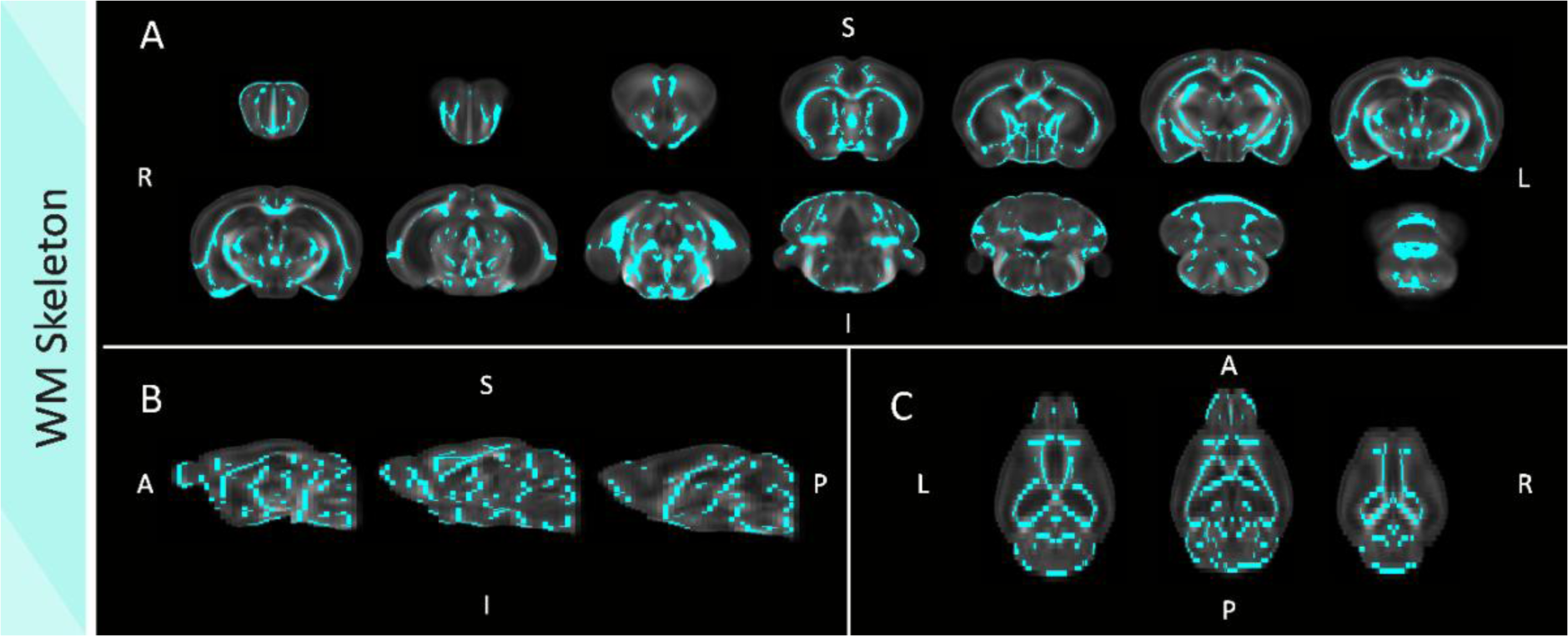
The MouseX DW-ALLEN white matter skeleton. White matter skeleton superimposed over the FA-mean template. A) coronal, B) sagittal and C) axial serial slices of the calculated white matter skeleton, over the FA template contrast.

Once the MRI based template was generated, we proceeded to its anatomical annotation. The MouseX DW-ALLEN annotation was created with the AMBA CCFv3 annotation as a starting point (Figure 4A). Since typical MRI preclinical image have much lower resolution compared to the AMBA CCFv3 template, AMBA annotation was re-structured and highly condensed for the creation of the MouseX DW-ALLEN annotation, merging regions whose resolution was too low. Our annotation contains a total of 63 regions of interest, respecting the anatomical hierarchy of the Allen Mouse Brain: 27 regions corresponding to the cortical plate, 3 to the cortical subplate, 6 cerebral nuclei, 8 brainstem regions, and the cerebellum conform the grey matter annotation; white matter annotation delineates 17 fiber tracts groups; and the ventricular system is individually delineated. Table 1 collects the 63 regions and the structure size, and Figure 4B, C, D, E and G show the atlas annotation. As a result, the MouseX DW-ALLEN annotation delineates the T2W template with high accuracy, detailing, and correspondence to the Allen Mouse Brain annotation (as shown in the close-up example in Figure 4F, G and H). To account for multi-contrast differences, white matter annotation was further corrected according to the DW-based templates FA and MD (Figure 4G (right) and H (right)).

**Figure 4.**
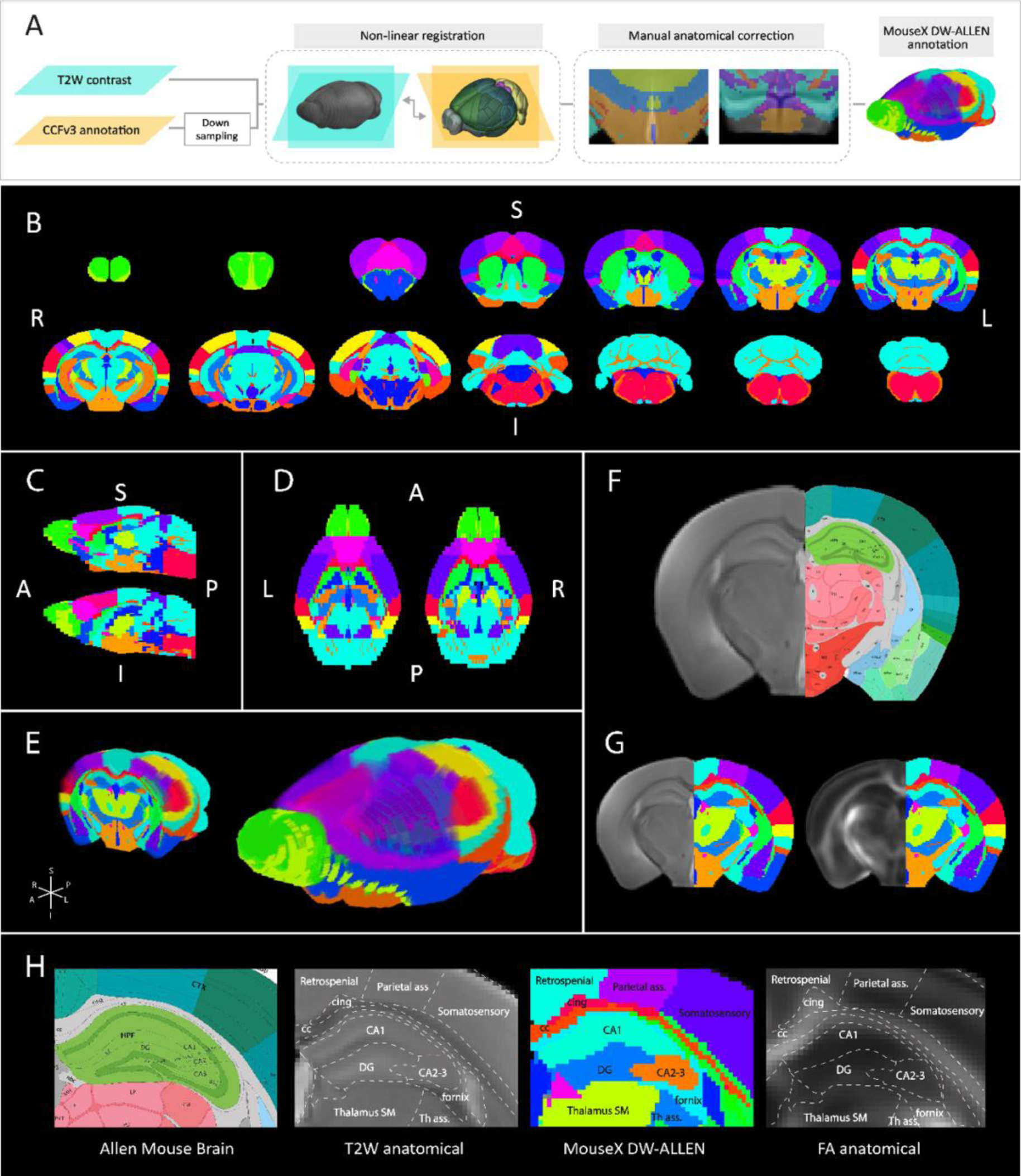
The MouseX DW-ALLEN Atlas. The MouseX DW-ALLEN annotation generation. A) Workflow of the generation of the annotation, taking CCFv3 annotation as a starting point, registered non-linearly to the MRI T2W template contrast, and manually and exhaustively corrected to anatomically match the template. B) Coronal, C) sagittal and D) axial serial slices of the MouseX DW-ALLEN annotation, and E) three-dimensional rendering. F) Accuracy of the anatomical annotation matching between Allen Mouse Brain Atlas annotation (right half) and G) the MouseX DW-ALLEN annotation, superimposed over a representative template T2W contrast slice (left) and FA contrasts (right). H) Close up visualization of the annotation details, comparing the hippocampal region and surrounding cortices and subcortical nuclei, in the Allen Mouse Brain reference Atlas, T2W template contrast, MouseX DW-ALLEN annotation and FA contrast.

**Table 1.**
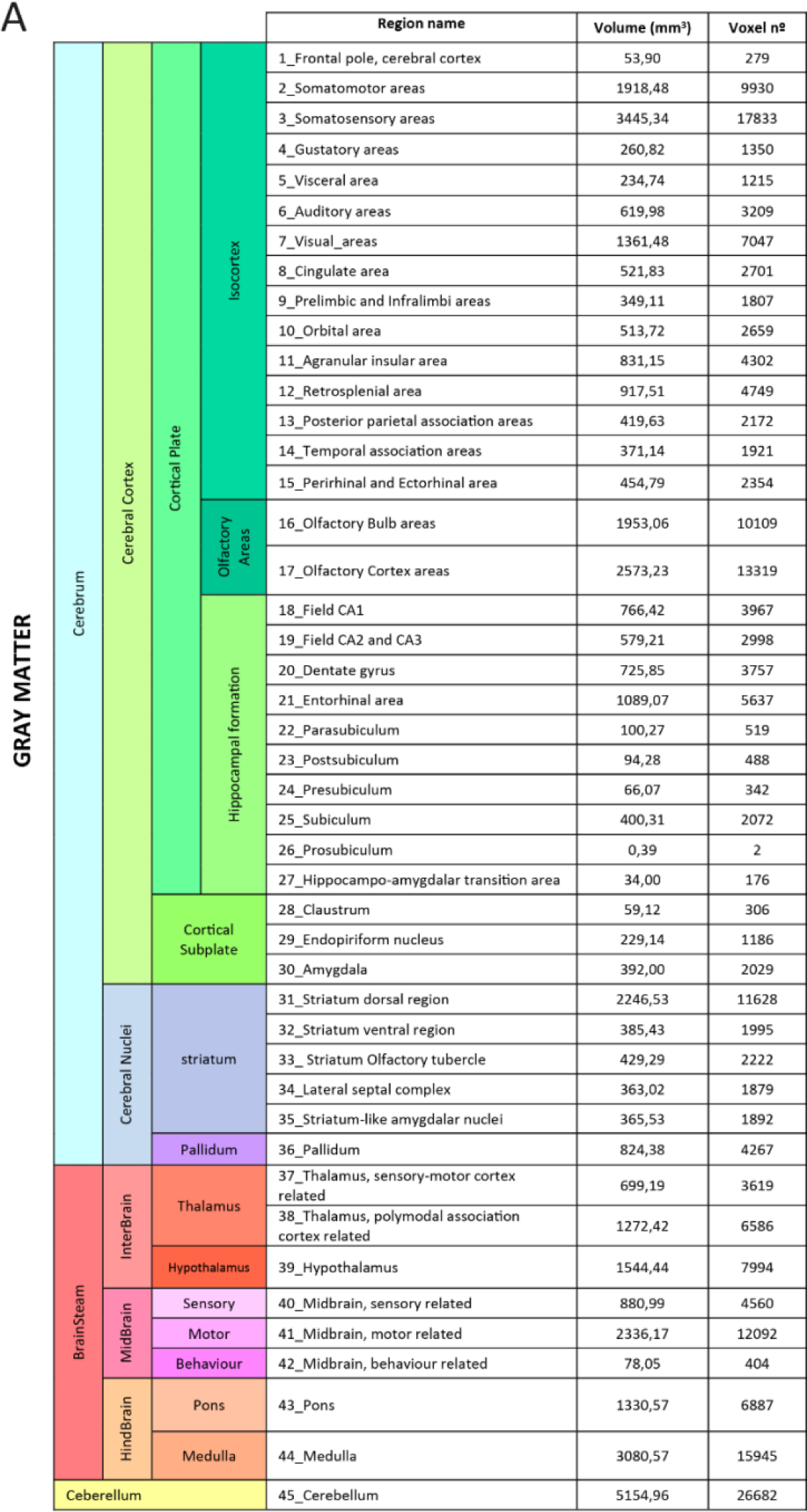
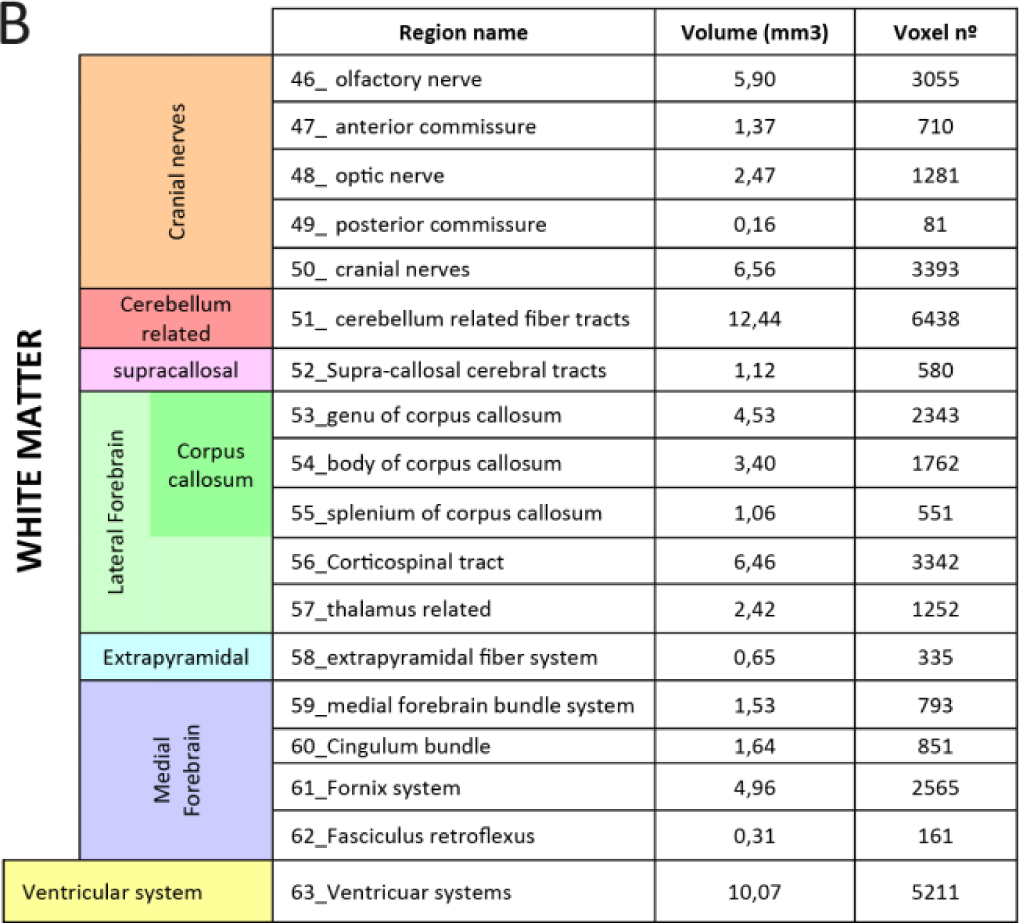
A) Gray matter and B) White matter annotation of the MouseX DW-ALLEN Atlas into regions and subregions, following anatomical divisions of Allen Mouse Brain Atlas. Regions are noted with their corresponding numeric code.

### Leveraging Allen Resources: mean diffusivity in grey matter is sensitive to microglia density

To demonstrate the importance of sharing a common annotation between different neuroimaging techniques, we correlated ROI-specific MD values with general cell count, and with each specific cell population present in the 3D Cell Atlas [20]: neurons, whole glial count, astrocytes, oligodendrocytes, and microglia. Correlation between cell count and MD signal was statistically significant only for microglia count (p-value= 0.041) (Figure 5 and Supplementary Figure 2) and not for the other cell types (astrocytes, p-value=0.24; oligodendrocytes, p-value=0.59; general glial count, p-value=0.93; neurons, p-value=0.96; and whole cell count, p-value=0.97).

**Figure 5.**
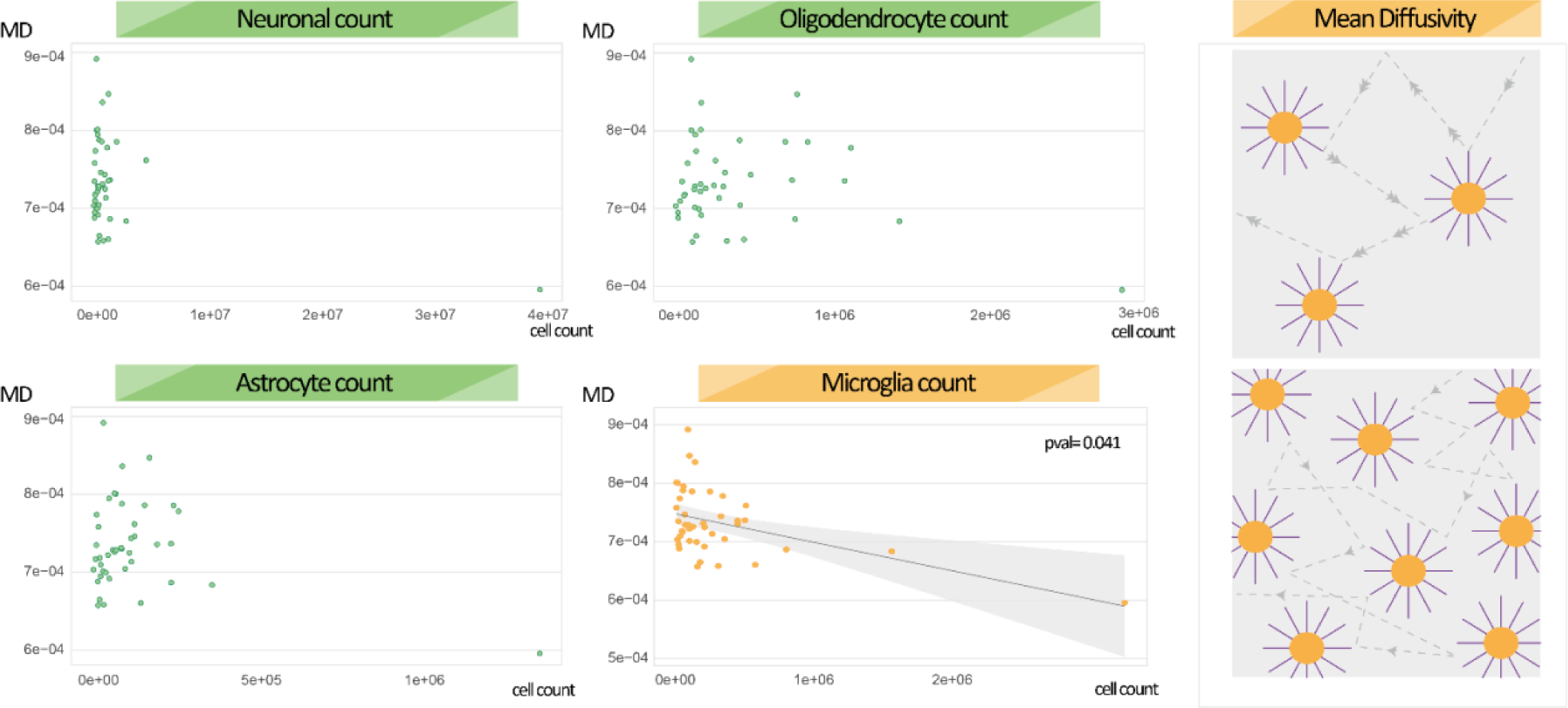
Microglia cell count correlates with mean diffusivity. Mean diffusivity is significantly associated with microglia count – and not other cell count – from [20]. Right panel: schematic illustration of water diffusion for low (upper figure) and high (lower panel) microglia densities.

### Sex differences: using a male-only template generates spurious sex effects

To quantify the bias introduced by the use of a male-based 3D template, we compared the deformation fields across sexes through voxel-wise statistics. Widespread significant positive and negative differences were evidenced (Figure 6A). On the other hand, when female subjects were included for the generation of the averaged 3D space, the deformation fields are not statistically different across sexes when matched to the template, except for a small region belonging to the olfactory cortex (Figure 6B). When the two templates are used on new mice longitudinal DW-MRI data to test for sex effects, the male-only template detects robust sex differences across multiple regions, which are however not detected when using our MouseX DW-ALLEN Atlas, built with both sexes. The analysis is reported in Figure 7.

**Figure 6.**
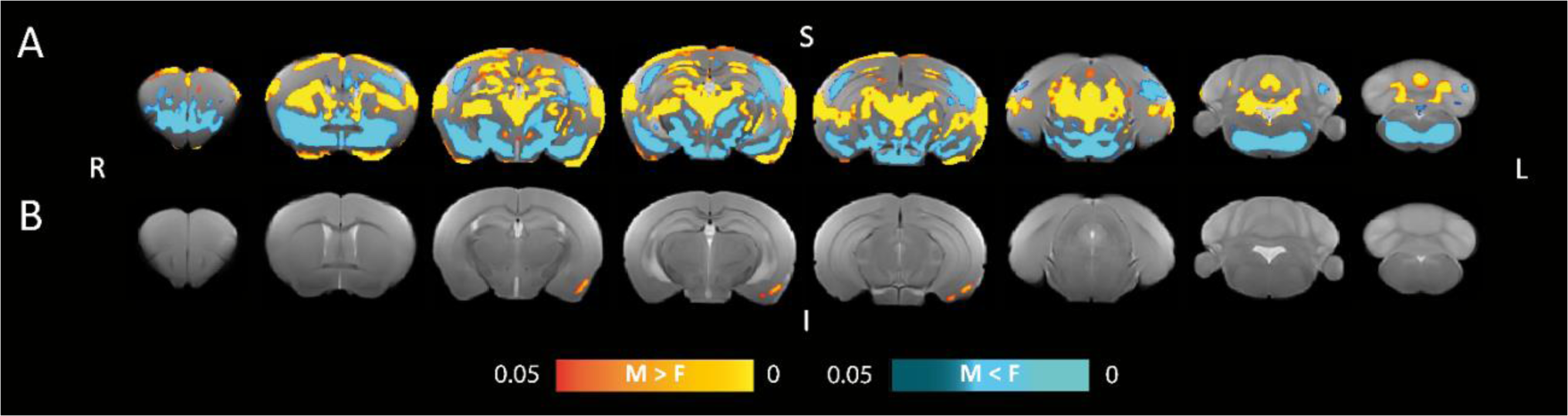
Comparison of the Jacobian across sexes. Voxel-wise comparison across sexes of the Jacobian of the transformation to match A) a male-only template and B) the MouseX DW-ALLEN Template, generated using animals of both sexes. The comparison when matching a female-only atlas is reported in Supplementary Figure 1. Yellow-red voxels indicate male Jacobian larger than female Jacobian, while blue-lightblue represent the opposite contrast.

**Figure 7.**
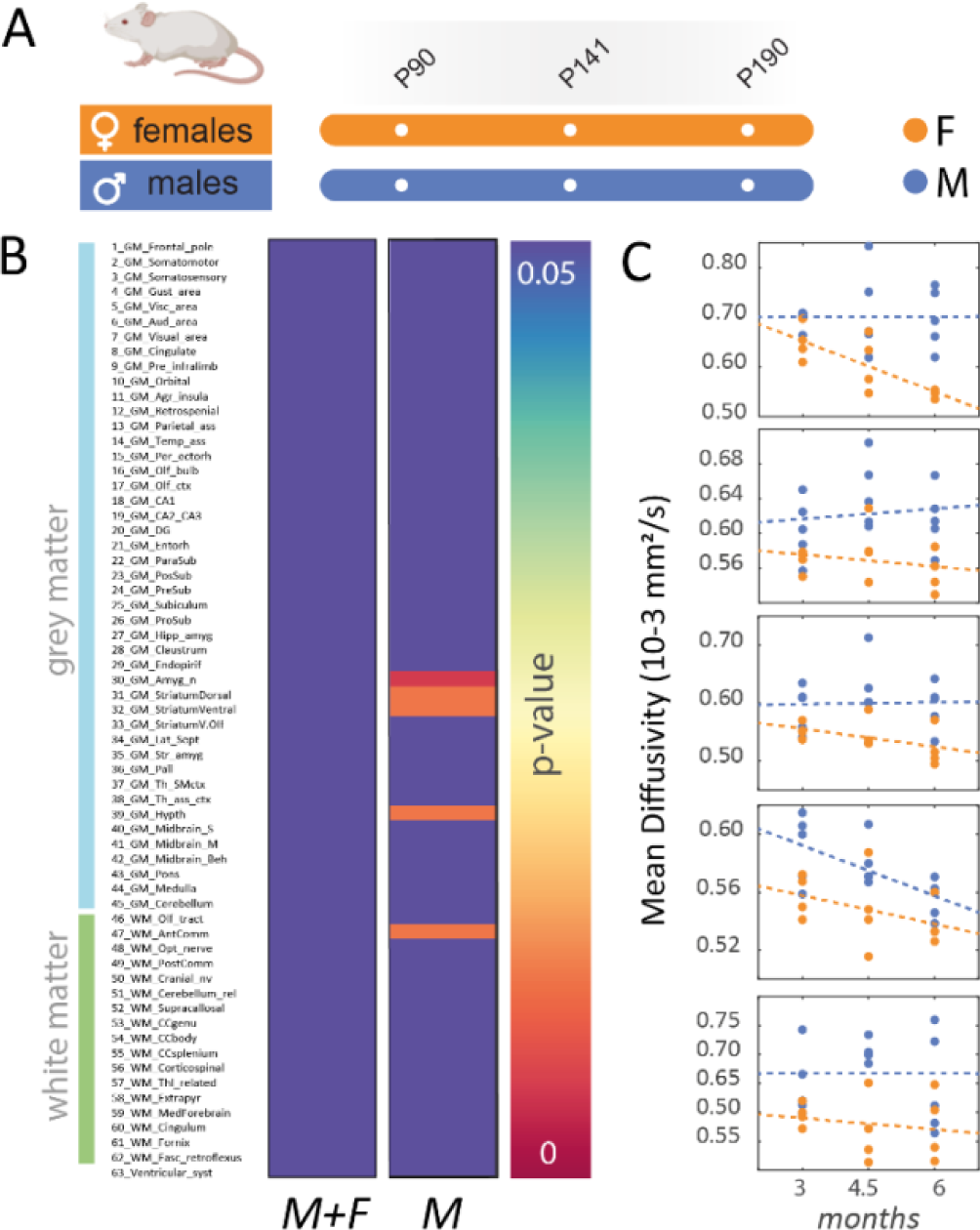
Using a male-only template generates spurious sex effects in a longitudinal mice experiment. A) Experimental design: 4 female and 5 male mice underwent a DW-MRI longitudinal experiment at 3, 4.5 and 6 months of age. B) A general linear model was used in each region to test for significant sex effects in two analyses: when using a template built with animals of both sexes (left) and male only (right). No significant sex effects are present in the first analysis, while 5 regions have significant sex effect in the second analysis. C) Regions where significant sex effect is detected when using a male-only template, respectively: amygdala, ventral and dorsal striatum, hypothalamus and anterior commissure.

## Discussion

Diffusion weighted MRI (DW-MRI) holds significant potential for addressing both clinical and preclinical research questions. As more advanced approaches are being developed, its capacity to disentangle the biological substrate in healthy and pathological brain parenchyma grows bigger. For example, diffusion weighted MRI (DW-MRI) has recently been proved capable to reveal neuroinflammation in different brain conditions [30], [31]. Recent efforts have used animal models to demonstrate that DW-MRI is sensitive to changes in glia morphology and proliferation [4]. This solidifies MRI’s standing as a highly effective tool in preclinical research. However, to express its full potential, DW-MRI methods should be also applicable in mice, to tap into the vast possibility of genetic modifications. This calls for specifically designed tools to aid the data analysis.

Standardized atlases are pivotal in the processing pipeline, being necessary for both ROI-based and voxel-wise statistical analyses. Given the existence of a consensus mouse brain atlas, the Allen Mouse Brain Atlas (AMBA), that has been widely adapted for many neuroimaging fields [17], [32], [33], and which contains a unique set of molecular and genetic resources, a desirable characteristic of the mouse MRI atlas would be to feature the same AMBA annotation. Another important desirable feature for a standardized atlas is the inclusion of animal of both sexes, considering the efforts to overcome the well-known gender bias. In addition, to achieve a generalizable resource, the atlas should be based on a large cohort of animals, possibly scanned *in vivo*. Indeed, perfusion and fixation alter the tissue diffusion properties, so that DW-MRI *in vivo* and *ex vivo* contrasts are inherently different [34].

Currently available mice MRI atlases are limited to a regional parcellation such as neocortex [35], to whole brain divisions no larger than of 40 regions [36]–[38], or to *ex vivo* MRI [39]. To date, the most exhaustively defined MRI whole brain mice atlas available was manually defined by Dorr et al in 2008 [40], which divides the brain into 62 regions, but is dedicated to *ex vivo* contrast and lacks the correspondence to the Allen database. Moreover, previous efforts of anatomical delineation have been done by generating a template based on a limited sample size [36], [41], [42], lacking thus sensibility to account microstructural differences between sexes.

To fill this gap, we generated the MouseX DW-ALLEN Atlas, the first *in vivo* DW-MRI based 3D mouse whole-brain resource sharing the AMBA delineation generated using a large cohort of animals of both sexes, and specific for DW-MRI contrast. Its parcellation into 63 regions has the optimal anatomical subdivision for common preclinical MRI resolution, and allows the precise region-wise dissection of MRI signal.

While single sex approaches have been predominant to date, there is solid evidence of sexual dimorphism in mice brain anatomy and function [18], [43]–[45]. Therefore, including both sexes in generating standardized resources is a necessary step to overcome the gender bias. We demonstrate that males and females undergo significantly different deformations when matching a single-sex template, in accordance with a recent Voxel Based Morphometry study [43]. Instead, when using a template generated with both sexes, the sexual dimorphism is compensated, and the difference in the warp field between males and females is substantially reduced. Importantly, the use of a male-only template, commonplace in preclinical research, generates a bias in the data analysis, so that spurious sex differences appear. These results have far-reaching implications that go beyond the present context, and will hopefully foster a change of paradigm towards a more balanced representation of females in research.

We finally aimed to demonstrate the huge potential of providing an atlas based on the commonly used AMBA annotation. The *In Situ Hibridization* (ISH) database from Allen brings together the expression data of approximately 20.000 distinct mouse genes [13], what allowed the construction of the Cell Atlas of the Mouse Brain [20] for cell type counting. As our data shares the common AMBA delineation, we demonstrated that region wise MD measures is negatively associated with microglia cell number, but not with other cell types. A previous study showed sensitivity of MD to neuroinflammation in the context of alcohol addiction [19]. In addition, in the same study, when microglia population is depleted with the CSF-1R inhibitor PLX5622, MD increases [19]. Thus, the negative correlation between microglia cell number and MD measures is in agreement with previous results. Interestingly, in the cerebellum both microglia cell-count and MD measures have the most extreme values. It is widely described in bibliography that the cerebellum possesses a unique microglial profile, with specific functional dynamics and phenotype [46]. Cerebellar microglia are less ramified and sparsely distributed [47]–[50], thus, are a plausible substrate for a lower MD.

### Limitations

This atlas holds, nevertheless, few limitations. For time constrains, the chosen voxel resolution is anisotropic in the axial plane. While suboptimal for tractography, this is commonplace in mice *in vivo* MR imaging.

In addition, correlation between MD and cell density should be carefully interpreted. MD has been proven to be sensitive to microglia, but it is not specific, which means that there are more components contributing to MD variability. Parameters more specific to microglia proliferation are expected to have stronger association with microglia count.

In summary, the MouseX DW-ALLEN Atlas fills an important gap in preclinical imaging research, promotes the inclusion of animals of both sexes in preclinical research and opens a promising scenario of collaborative and multi-approach science.

## Author contributions

Conceptualization: PMT and SDS; Data curation: PMT, AS and SDS; Formal analysis: PMT, AS and SDS; Funding acquisition: SDS; Investigation: PMT, AS and SDS; Methodology: PMT, EES, MKS and SDS; Project administration: SDS; Resources: SDS; Software: MKS and SDS; Supervision: SDS; Validation: SDS; Visualization: PMT and SDS; Writing: PMT and SDS.

## Acknowledgements

We thank Aroa Sanz Maroto for excellent technical support. We also thank the *In Vivo* Imaging facility of the Instituto de Neurociencias for data acquisition.

