## Supplementary Figures for "Sex matters: the MouseX DW-ALLEN Atlas for mice diffusion-weighted MR imaging"

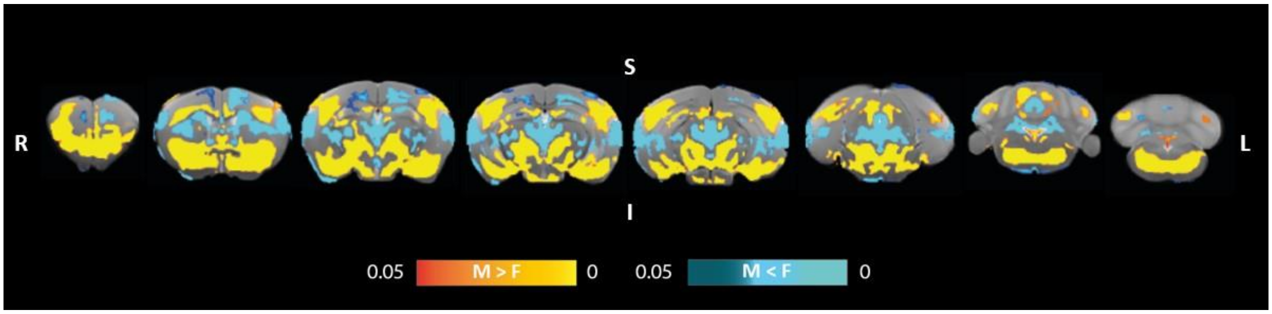


**Supplementary figure 1. Comparison of the Jacobian for female-only atlas.** Voxel-wise comparison across sexes of the Jacobian of the transformation to match a female-only atlas. Yellow-red voxels indicate male Jacobian bigger than female Jacobian, while blue-lightblue represent the opposite contrast.

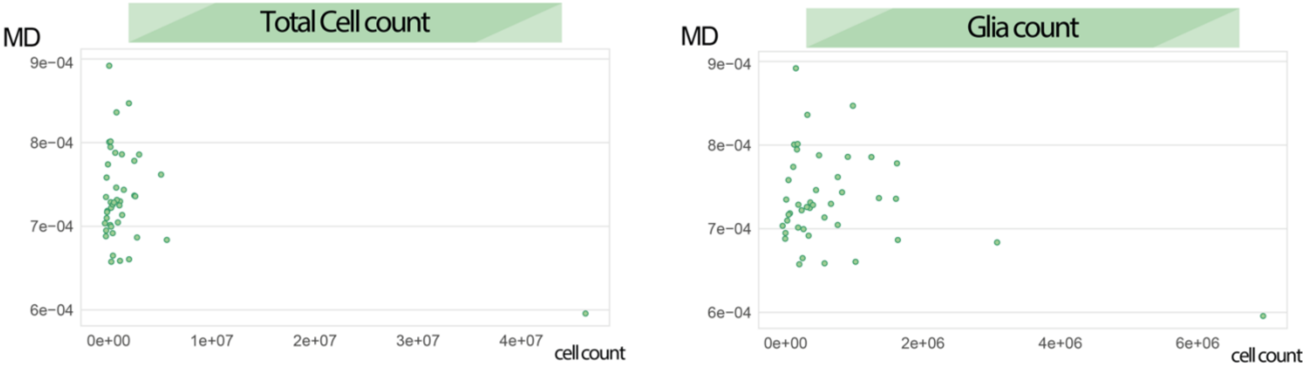


**Supplementary figure 2. Association between cell counts and mean diffusivity.** Scatterplot of total cell count (left) and total glia count (right) from [20] versus mean diffusivity.
